# Low-Dose Microcystin-LR Elicits Sex-Dimorphic Transcriptomic Responses in Senescent *Nothobranchius furzeri*: Implications for Cyanotoxin Vulnerability in Aging Vertebrates

**DOI:** 10.64898/2026.08.17.745368

**Authors:** Zainab Afzal, Charles Hatcher, Deepak Kumar

## Abstract

Microcystin-LR (MC-LR), a cyanobacterial toxin produced during harmful algal blooms, is an increasing environmental and public health concern as the frequency and intensity of harmful algal blooms continue to rise globally. While the effects of MC-LR have been extensively studied in young organisms, much less is known about how aging influences susceptibility to cyanotoxin exposure. Here, we used the naturally short-lived turquoise killifish, *Nothobranchius furzeri*, to investigate transcriptional responses to low-level MC-LR exposure in a senescent vertebrate. Approximately 8-month-old GRZ killifish were exposed to a low dose of 0.5 µg/L MC-LR, followed by whole-body RNA sequencing and sex-stratified differential expression analysis. Despite identical experimental conditions and exposure, males and females exhibited strikingly distinct transcriptional responses, with 313 differentially expressed genes (DEGs) in males and 263 in females and only 27 DEGs shared between the sexes. Among the shared responses, *pck1*, a key regulator of gluconeogenesis, was strongly downregulated in both sexes, accompanied by altered expression of genes associated with mitochondrial function, metabolic regulation, extracellular matrix remodeling, and genome maintenance. Males exhibited prominent remodeling of skeletal muscle and contractile programs, supported by enrichment of sarcomeric, myofilament, and contractile-fiber-associated genes. In contrast, females showed pronounced alterations in reproductive and metabolic programs, including vitellogenin- and zona pellucida-associated transcripts. Cell/tissue associated marker-module analysis further revealed distinct sex-dependent shifts in structural, neural, immune, metabolic, and reproductive transcriptional signatures. Together, these findings demonstrate that MC-LR elicits a broad but strongly sex-dependent transcriptional response in senescent *N. furzeri*, involving responses in multiple physiological systems. Our study identifies biological sex as an important determinant of cyanotoxin responses in an aging context and establishes naturally aged *N. furzeri* as a tractable vertebrate model for investigating interactions between environmental exposure and biological aging.

## Introduction

The African turquoise killifish, *Nothobranchius furzeri*, provides a unique vertebrate model for investigating how biological aging influences susceptibility to environmental stressors. The short-lived GRZ strain undergoes rapid and well-characterized aging including curved spines (lordosis), frayed fins, faded or lost coloration, progressive declines in locomotor activity characterized by sluggish movement and reduced swimming capacity, along with deterioration in tissue homeostasis, regenerative capacity, and other molecular and physiological hallmarks associated with vertebrate aging (Bedbrook et al., 2023; Harel et al., 2015; Hu & Brunet, 2018; Ma et al., 2025; Otsuka & Matsui, 2025; Polacik et al., 2016; Valdesalici & Cellerino, 2003; Valenzano et al., 2006). Under laboratory conditions, *N. furzeri* has a remarkably compressed lifespan, with the GRZ strain typically reaching a median lifespan of approximately 3–7 months. By approximately 8 months of age, these fish representing a naturally senescent cohort, correspond broadly to an advanced human age of 90+ years when considered in the context of their highly compressed lifespan (Harel et al., 2016). Furthermore, the highly inbred nature of the GRZ strain (Valenzano et al., 2009) provides genetically homogeneous cohorts, reducing inter-individual genetic variability and offering an important advantage for resolving age- and exposure-associated molecular responses in transcriptomic studies. Together, the compressed lifespan and genetic homogeneity make the GRZ strain a powerful model for investigating environmental exposures within the physiological context of advanced biological aging while maintaining experimentally tractable timescales.

One environmental exposure of growing concern is the cyanobacterial harmful algal blooms (CyanoHABs), which are increasing in frequency, duration, and geographic distribution as rising global temperatures and anthropogenic nutrient enrichment from agricultural runoff and urbanization are creating conditions that favor bloom formation (Feng et al., 2024). CyanoHABs can produce microcystins (MCs), a diverse family of cyclic peptide toxins associated with increasing ecological and human health concerns (McLellan & Manderville, 2017). More than 280 MC types have been identified, with human exposure primarily occurring through contaminated drinking water and food, recreational contact with affected waters, and consumption of contaminated algal dietary supplements (Bouaicha et al., 2019; Bruno et al., 2006; Carmichael et al., 2001; Carmichael et al., 1988; Carmichael & Bent, 1981; Gilroy et al., 2000; Gorham, 1962; Gupta et al., 2024; Kappers et al., 1981; Vinogradova et al., 2011; Wang et al., 2023; Watanabe et al., 1988). Among the different MC types, microcystin-LR (MC-LR) is one of the most extensively studied and it is a potent inhibitor of the serine/threonine protein phosphatases PP1 and PP2A. Following uptake through organic anion transporting polypeptides, particularly in the liver, MC-LR can disrupt cellular phosphorylation (disrupting the phosphorylation state of hundreds of downstream signaling proteins), cytoskeletal organization, oxidative homeostasis, mitochondrial function, and energy metabolism (Diez-Quijada et al., 2019; Duan et al., 2025; Li et al., 2025; Maric et al., 2021; Martins et al., 2017; Meng et al., 2011; Rajpoot et al., 2025; Soni et al., 2026). Despite this well-established toxicity, we know considerably less about how these molecular responses are altered when exposure occurs in the physiological context of advanced biological aging.

This knowledge gap is particularly important because aging progressively alters many of the same cellular systems targeted by environmental toxicants. Older organisms exhibit changes in mitochondrial function, proteostasis, DNA damage responses, immune regulation, and metabolic homeostasis (Lopez-Otin et al., 2023), potentially reducing their capacity to buffer additional environmental stress. Yet, much of the experimental literature defining MC-LR toxicity has relied on young or reproductively mature organisms (Breidenbach et al., 2025; Liu et al., 2018; Lone et al., 2015; Ma et al., 2026; Wang et al., 2023), cell-based models (Ikumawoyi et al., 2024; Li et al., 2011; Niture et al., 2025), or organoids (Chowdhury et al., 2024; Y. Liu et al., 2026), leaving the effects of cyanotoxin exposure in aged organisms comparatively understudied. Hence, concentrations that produce limited effects in younger organisms may elicit mechanistically different and more severe molecular responses when exposure occurs against the altered physiological background of aging. Understanding this interaction is increasingly relevant because aging populations and the growing prevalence of harmful algal blooms increase the potential intersection between biological aging and cyanotoxin exposure.

Biological sex may further modify this response. Male and female vertebrates differ in endocrine signaling, hepatic metabolism, reproductive energetic demands, and xenobiotic processing (Della Torre & Maggi, 2017; Oliva et al., 2020; Zheng et al., 2018), all of which can influence toxicant pharmacokinetics and shape the magnitude and nature of transcriptional responses to environmental contaminants. These differences may become particularly important with aging, as endocrine and metabolic states shift from those observed earlier in life. It remains largely unresolved how biological sex and aging shape the molecular response to cyanotoxin exposure, particularly at the transcriptomic level. Examining males and females independently within a senescent population, where endocrine and physiological states have shifted considerably from reproductive-age baselines, may reveal sex-specific molecular responses that would otherwise be obscured when sex is treated solely as a covariate or when both sexes are analyzed together.

Here, we used approximately 8-month-old *N. furzeri* GRZ adults to investigate sex-specific transcriptional responses to low-dose MC-LR exposure in an aged vertebrate. Fish were exposed to 0.5 µg/L MC-LR, a concentration selected within the range of health-based exposure guidance values (*Toxic Cyanobacteria in Water*, 2021), followed by whole-body transcriptomic profiling with sex-stratified analyses. We identify both shared and strongly sex-divergent molecular responses, including disruption of metabolic and mitochondrial-associated programs together with distinct muscle- and reproductive-associated transcriptional signatures in males and females. These findings establish biological age and sex as important contexts for understanding cyanotoxin responses and demonstrate the utility of the short-lived, naturally senescent, killifish for investigating how environmental exposures intersect with the biology of aging.

## Materials and Methods

### Animal husbandry

Adult African turquoise killifish (*Nothobranchius furzeri*, GRZ strain), a highly inbred strain, were maintained in 2.78-L tanks within a recirculating aquatic system (Aquaneering, San Diego, CA, USA) containing reverse osmosis (RO) system water at 26–27 °C under a 14:10 h light:dark cycle. Water pH was maintained between 7.0 and 7.5. Fish were fed live brine shrimp (EG Artemia, INVE Aquaculture Inc.) three times daily and supplemented with Otohime fish diet in (Reed Mariculture, Otohime C1) and bloodworms (Hikari Bio-Pure Blood Worm Cubes; That Fish Place, Catalog No. 284799). Males and females were individually housed, and sex was determined based on secondary sexual characteristics, including differences in coloration and body size. All killifish husbandry and experimental procedures followed established protocols (Nath et al., 2023), and were approved by the Institutional Animal Care and Use Committee (IACUC) at North Carolina Central University (Protocol No. DK10112024).

### Chemical exposure

Senescent adult killifish (∼8 months old) were removed from the recirculating system and acutely exposed to microcystin-LR (MC-LR; Cat. No. ALX-350-012-C100) at a concentration of 0.5 µg/L. MC-LR was diluted in reverse osmosis (RO) water, and individual fish were maintained in stand-alone 2.8-L containers containing either MC-LR treatment solution or RO water alone for 48 h. Exposure and control water were replaced daily to maintain consistent experimental conditions. Following the 48-h exposure, fish were euthanized by tricaine overdose and maintained in the euthanasia solution for approximately 20 min until complete cessation of opercular movement was observed. Whole bodies were then immediately flash-frozen in liquid nitrogen and stored at −80 °C until further processing.

### RNA extraction and sequencing

Whole-body homogenates were prepared from flash-frozen fish (n = 2 males and 2 females per treatment group; 8 total samples (K1-K8). Each sample was homogenized in 1mL TRIzol Reagent (Invitrogen, Cat. No. 15596018), and total RNA was isolated using the Direct-zol RNA Miniprep Kit (Zymo Research, Cat. No. R2073) according to the manufacturer’s instructions, including on-column DNase treatment. RNA concentration and purity were assessed using a NanoDrop One spectrophotometer (Thermo Fisher Scientific), and all samples exhibited A260/A280 ratios greater than 1.8. RNA samples were submitted to Novogene (Sacramento, CA, USA) for library preparation and sequencing. Libraries were generated using the Illumina TruSeq Stranded mRNA Library Preparation Kit and sequenced on the Illumina NovaSeq platform to produce 150-bp paired-end reads. All 8 samples passed pre-defined sequencing quality thresholds: Q30≥97.1%, error rate 0.01%, clean read retention 96–99%.

### Differential expression analysis

Clean reads were aligned to the Nothobranchius furzeri reference genome (Nfu_20140520; NCBI accession GCF_001465895.1) using HISAT2. Differential gene expression analysis was performed in R using the edgeR package. Lowly expressed genes were filtered prior to normalization, and count data were normalized using the trimmed mean of M-values (TMM) method. Gene-wise dispersion estimates were calculated using the negative binomial framework implemented in edgeR, and differential expression was assessed using generalized linear models. Resulting p-values were adjusted for multiple hypothesis testing using the Benjamini–Hochberg false discovery rate (FDR) correction. Genes with an FDR-adjusted p-value ≤ 0.05 and an absolute log₂ fold change (|log₂FC|) ≥ 2.0 were considered significantly differentially expressed. Three planned comparisons were performed to evaluate overall and sex-specific responses to MC-LR exposure: (1) all MC-LR-treated fish versus all untreated controls (K5–K8 vs. K1–K4); (2) MC-LR-treated males versus untreated males (K5–K6 vs. K1–K2); and (3) MC-LR-treated females versus untreated females (K7–K8 vs. K3–K4). Based on the direction of the contrasts used in the analysis, positive log₂FC values indicate higher expression in untreated controls and therefore decreased expression following MC-LR exposure, whereas negative log₂FC values indicate higher expression in MC-LR-treated fish and therefore increased expression following exposure.

### Functional enrichment and ortholog annotation

Statistical analyses and data visualization were performed in R. Gene Ontology (GO) and Kyoto Encyclopedia of Genes and Genomes (KEGG) pathway enrichment analyses were conducted using the clusterProfiler package. Over-representation analysis was performed using a hypergeometric test, with p-values adjusted for multiple comparisons using the Benjamini– Hochberg method. GO terms and KEGG pathways with an adjusted p-value ≤ 0.05 were considered significantly enriched. Overlap between differentially expressed genes identified in the male and female sex-stratified comparisons was determined by intersection of gene identifiers. Human orthologs for selected *N. furzeri* DEGs were identified using Swiss-Prot annotations provided in the Novogene gene-description output and cross-referenced against UniProtKB. Where direct orthology could not be established, genes were reported as putative functional homologs based on protein sequence similarity rather than definitive one-to-one human orthologs.

### Principal component analysis of global transcriptomic profiles

Principal component analysis (PCA) was performed to assess global transcriptomic variation among untreated and MC-LR-exposed senescent Nothobranchius furzeri. Gene-level expression values from the eight individual samples, comprising two biological replicates for each sex and treatment condition, were used for dimensionality reduction. Expression values were transformed prior to PCA to reduce the influence of highly expressed genes, and genes with insufficient or invariant expression across samples were excluded. PCA was performed across the complete retained transcriptome using the prcomp function in R, with samples represented according to treatment and sex. The proportion of total variance explained by each principal component was calculated from the corresponding eigenvalues and displayed on the PCA axes.

### Cell/tissue-associated transcriptional signature analysis

To examine whether MC-LR exposure was associated with coordinated changes in transcriptional programs representative of major cell and tissue types, a marker-based signature analysis was performed using the bulk RNA-sequencing expression profiles. Curated marker-gene sets representing neuronal, glial, skeletal muscle, fibroblast/extracellular matrix, endothelial, erythroid, hepatocyte, kidney, macrophage, neutrophil, T-cell, germ-cell, and proliferative transcriptional programs, based on published datasets (Teefy et al., 2023), were evaluated across individual samples. For each sample, a marker-module score was calculated for each signature by summarizing the normalized expression of genes assigned to the corresponding marker set. To account for the pronounced sex-dependent structure of the transcriptomic data, treatment-associated changes were calculated separately for males and females. For each transcriptional signature, module scores were averaged across biological replicates within each sex and treatment group, and the MC-LR-associated change was calculated as:

Δ module score= MC-LR module score −WT module score

Positive Δ module scores therefore indicate increased expression of the corresponding cell/tissue-associated transcriptional program following MC-LR exposure, whereas negative values indicate decreased expression relative to sex-matched untreated controls. These scores represent relative changes in marker-associated transcriptional programs and were not interpreted as direct estimates of cell-type abundance.

## Results

### Acute MC-LR exposure elicits sex-dependent transcriptional responses in aged killifish

To determine how an aged vertebrate responds to acute cyanotoxin exposure, approximately 8-month-old *Nothobranchius furzeri* (GRZ strain) males and females were exposed to 0.5 µg/L MC-LR for 48 h, followed by whole-body RNA sequencing to capture systemic transcriptional responses (Fig. 1A,B). All eight samples generated high-quality sequencing data, with Q30 scores ≥97.1% and 96–99% of reads retained following quality filtering.

**Figure 1.**
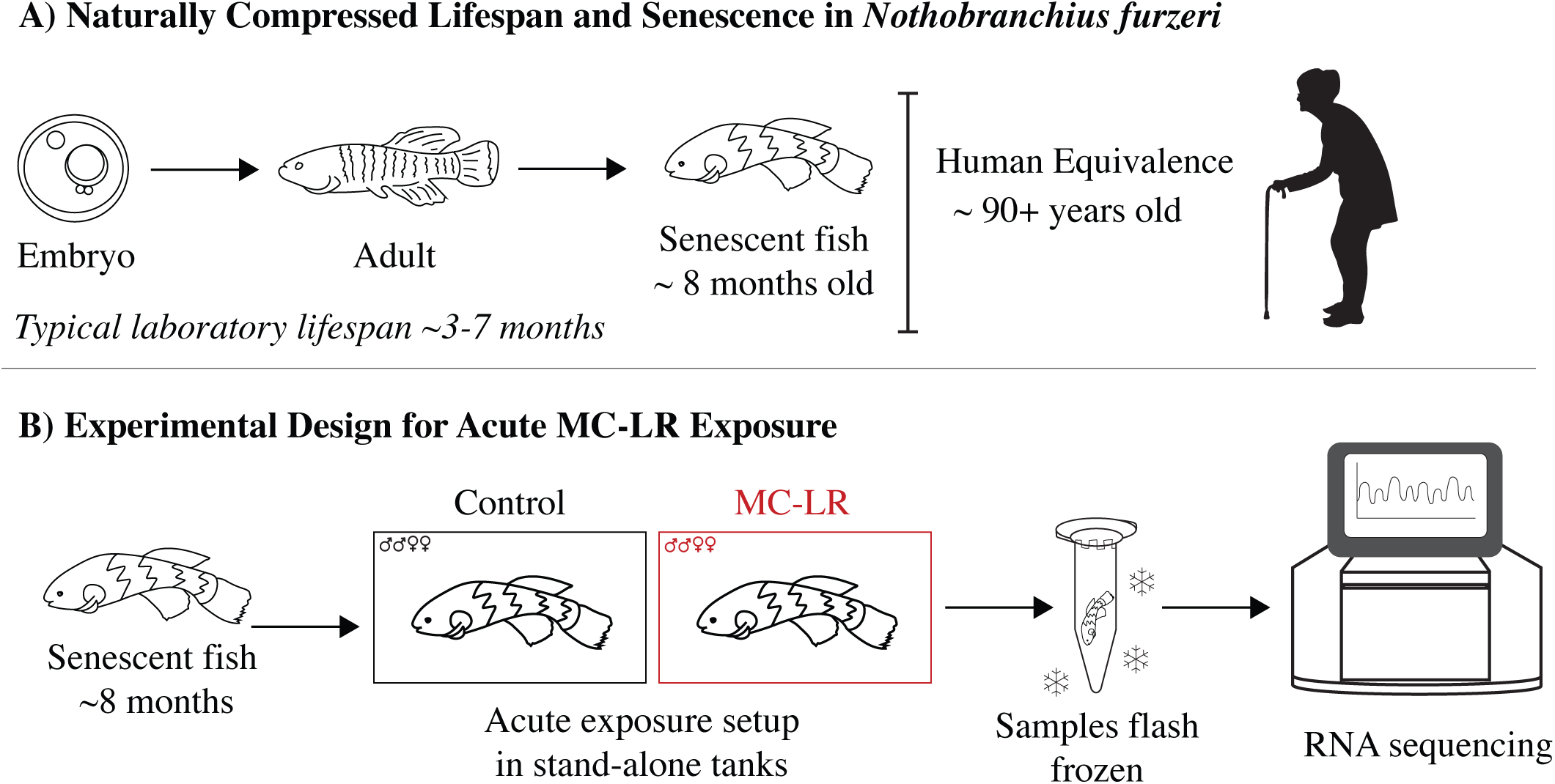
Senescent killifish model and experimental design for acute MC-LR exposure. (A) Schematic illustrating the naturally compressed lifespan of Nothobranchius furzeri. Under laboratory conditions, adults typically have a lifespan of approximately 3–7 months; thus, ∼8-month-old fish represent a naturally senescent population, broadly corresponding to an advanced human age of 90+ years based on comparative lifespan scaling. (B) Experimental design for acute MC-LR exposure. Senescent adult killifish (∼8 months old; two males and two females per condition) were exposed to 0.5 µg/L MC-LR for 48 h. Following exposure, whole-body samples were flash-frozen in liquid nitrogen and processed for RNA isolation. Samples were subsequently submitted to Novogene for library preparation and RNA sequencing.

We first assessed the transcriptional response to MC-LR across the entire cohort, irrespective of sex. This analysis identified 36 differentially expressed genes (DEGs), with 18 showing increased and 18 showing decreased expression following exposure (Fig. 2, left; Table 1, Supp Fig. 1). Sex-stratified analysis, however, revealed substantially broader responses. In males, MC-LR altered 313 genes, including 205 upregulated and 108 downregulated DEGs (Fig. 2, middle, Table 2, Supp Fig. 2). Females exhibited 263 DEGs, comprising 148 upregulated and 115 downregulated genes (Fig. 2, right; Table 3, Supp Fig. 3).

**Figure 2.**
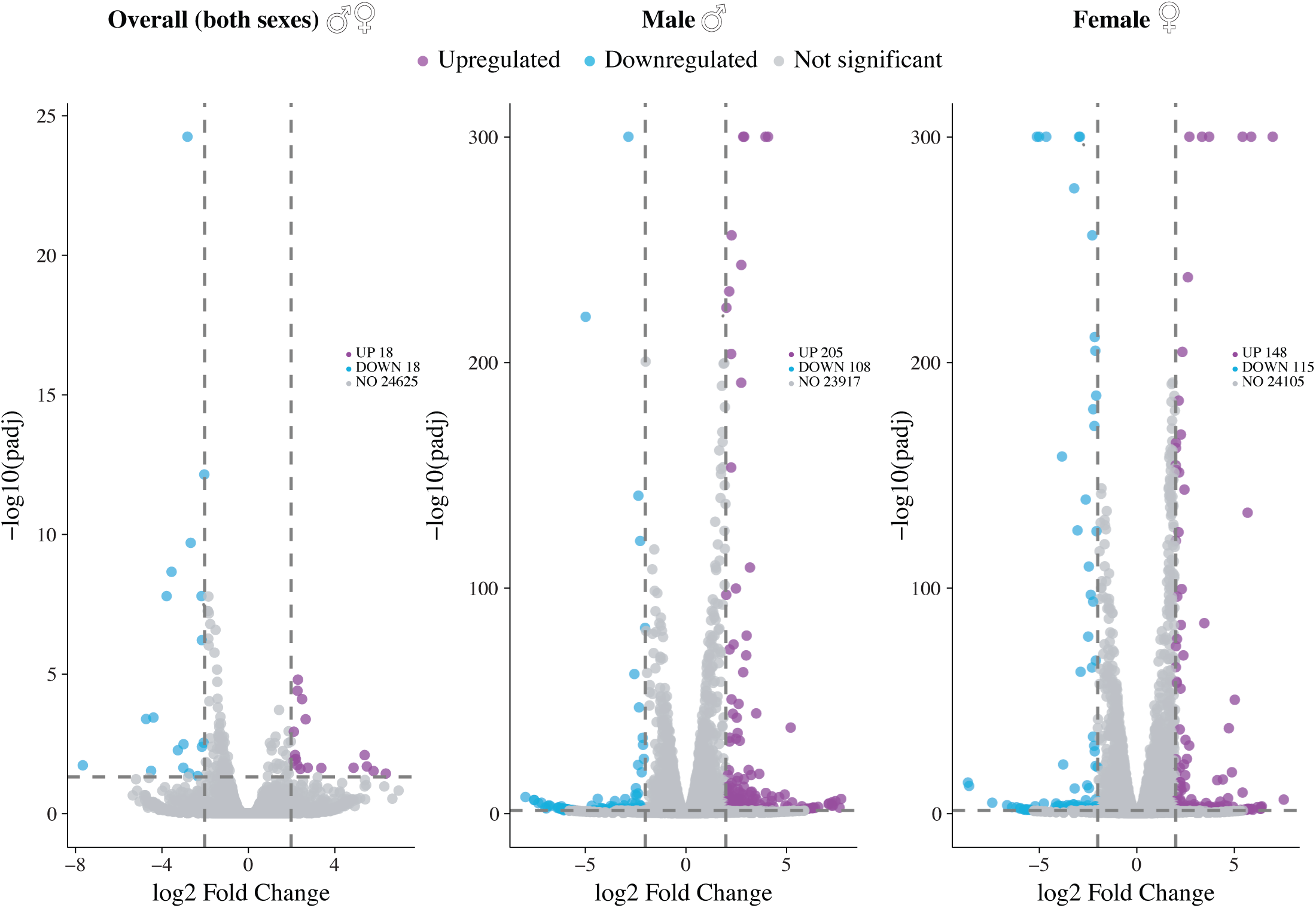
Differential gene expression in senescent Nothobranchius furzeri following MC-LR exposure. Volcano plots showing transcriptional responses to MC-LR exposure in the combined cohort (both sexes; left), males (middle), and females (right). Differential expression was assessed between MC-LR-exposed and untreated animals. Genes were considered significantly differentially expressed at an adjusted P value (padj) ≤ 0.05 and |log₂ fold change| ≥ 2. Purple points indicate genes upregulated following MC-LR exposure, blue points indicate downregulated genes, and grey points indicate genes not meeting the differential-expression thresholds. Dashed vertical lines denote log₂ fold-change thresholds of ±2, and the horizontal dashed line denotes the adjusted P-value significance threshold.

**Table 1:**
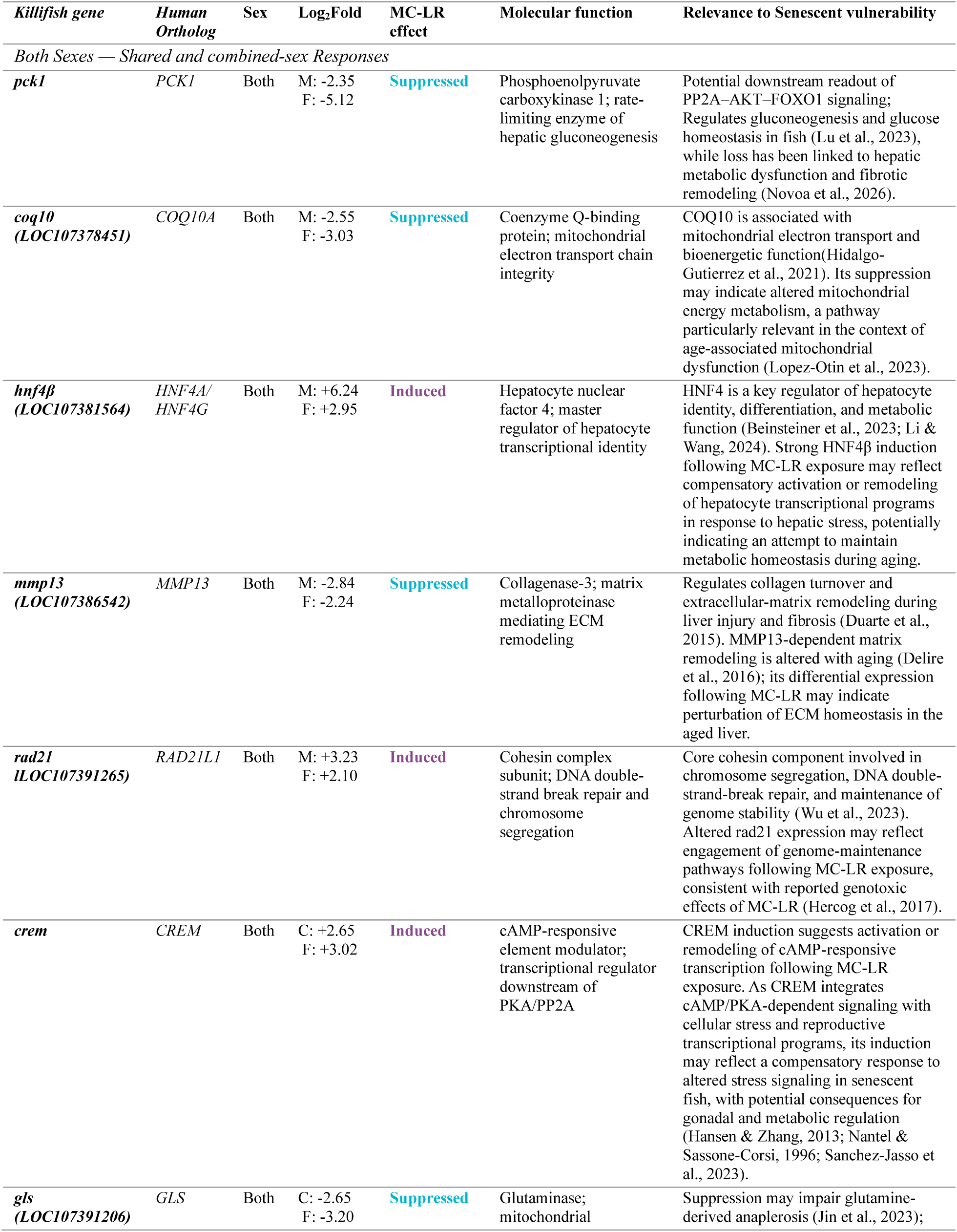
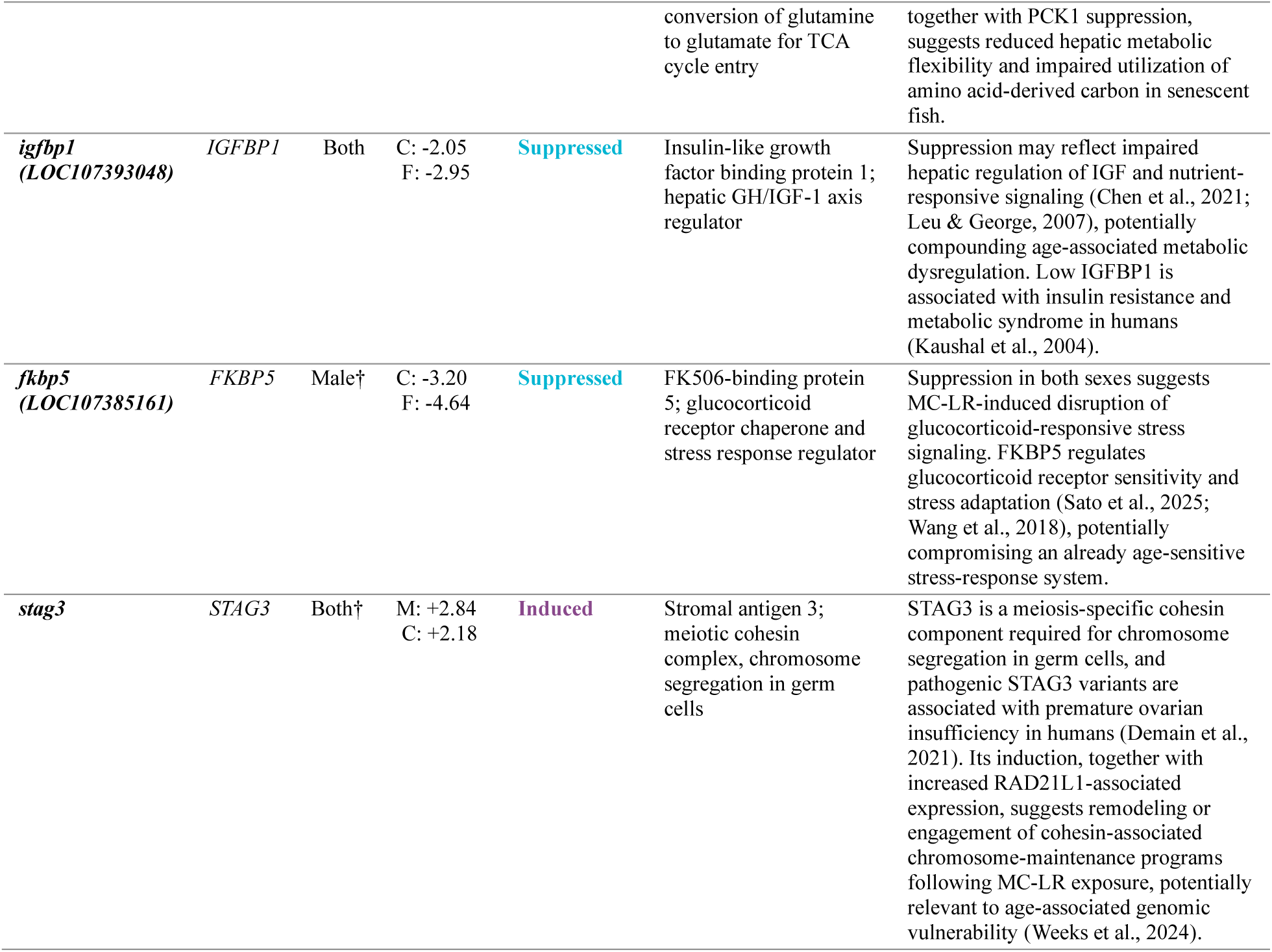
Selected differentially expressed genes in both sexes; log₂FC, log₂ fold change (positive values indicate induction by MC-LR; negative values indicate suppression by MC-LR). Human orthologs were inferred from Swiss-Prot annotations (Ahmad et al., 2025) and represent putative functional homologs based on protein sequence similarity rather than experimentally validated orthology. Selected differentially expressed genes in both sexes; log₂FC, log₂ fold change (positive values indicate suppression by MC-LR; negative values indicate induction by MC-LR). M: reflects log2fold in males, F: reflects log2fold in females, and C: reflect log2fold in combined analysis for both.

| *Killifish gene* | *Human Ortholog* | Sex | Log<sub>2</sub>Fold | MC-LR effect | Molecular function | Relevance to Senescent vulnerability |
| --- | --- | --- | --- | --- | --- | --- |
| *Both Sexes — Shared and combined-sex Responses* |  |  |  |  |  |  |
| *pck1* | *PCK1* | Both | M: -2.35<br>F: -5.12 | Suppressed | Phosphoenolpyruvate carboxykinase 1; rate-limiting enzyme of hepatic gluconeogenesis | Potential downstream readout of PP2A–AKT–FOXO1 signaling; Regulates gluconeogenesis and glucose homeostasis in fish (Lu et al., 2023), while loss has been linked to hepatic metabolic dysfunction and fibrotic remodeling (Novoa et al., 2026). |
| *coq10*<br>(*LOC107378451*) | *COQ10A* | Both | M: -2.55<br>F: -3.03 | Suppressed | Coenzyme Q-binding protein; mitochondrial electron transport chain integrity | COQ10 is associated with mitochondrial electron transport and bioenergetic function (Hidalgo-Gutierrez et al., 2021). Its suppression may indicate altered mitochondrial energy metabolism, a pathway particularly relevant in the context of age-associated mitochondrial dysfunction (Lopez-Otin et al., 2023). |
| *hnf4β*<br>(*LOC107381564*) | *HNF4A*/<br>*HNF4G* | Both | M: +6.24<br>F: +2.95 | Induced | Hepatocyte nuclear factor 4; master regulator of hepatocyte transcriptional identity | HNF4 is a key regulator of hepatocyte identity, differentiation, and metabolic function (Beinstainer et al., 2023; Li & Wang, 2024). Strong HNF4β induction following MC-LR exposure may reflect compensatory activation or remodeling of hepatocyte transcriptional programs in response to hepatic stress, potentially indicating an attempt to maintain metabolic homeostasis during aging. |
| *mmp13*<br>(*LOC107386542*) | *MMP13* | Both | M: -2.84<br>F: -2.24 | Suppressed | Collagenase-3; matrix metalloproteinase mediating ECM remodeling | Regulates collagen turnover and extracellular-matrix remodeling during liver injury and fibrosis (Duarte et al., 2015). MMP13-dependent matrix remodeling is altered with aging (Delire et al., 2016); its differential expression following MC-LR may indicate perturbation of ECM homeostasis in the aged liver. |
| *rad21*<br>(*LOC107391265*) | *RAD21L1* | Both | M: +3.23<br>F: +2.10 | Induced | Cohesin complex subunit; DNA double-strand break repair and chromosome segregation | Core cohesin component involved in chromosome segregation, DNA double-strand-break repair, and maintenance of genome stability (Wu et al., 2023). Altered rad21 expression may reflect engagement of genome-maintenance pathways following MC-LR exposure, consistent with reported genotoxic effects of MC-LR (Hercog et al., 2017). |
| *crem* | *CREM* | Both | C: +2.65<br>F: +3.02 | Induced | cAMP-responsive element modulator; transcriptional regulator downstream of PKA/PP2A | CREM induction suggests activation or remodeling of cAMP-responsive transcription following MC-LR exposure. As CREM integrates cAMP/PKA-dependent signaling with cellular stress and reproductive transcriptional programs, its induction may reflect a compensatory response to altered stress signaling in senescent fish, with potential consequences for gonadal and metabolic regulation (Hansen & Zhang, 2013; Nantel & Sassone-Corsi, 1996; Sanchez-Jasso et al., 2023). |
| *gls*<br>(*LOC107391206*) | *GLS* | Both | C: -2.65<br>F: -3.20 | Suppressed | Glutaminase; mitochondrial | Suppression may impair glutamine-derived anaplerosis (Jin et al., 2023); |
|  |  |  |  |  | conversion of glutamine to glutamate for TCA cycle entry | together with PCK1 suppression, suggests reduced hepatic metabolic flexibility and impaired utilization of amino acid-derived carbon in senescent fish. |
| *igfbp1*<br>(*LOC107393048*) | *IGFBP1* | Both | C: -2.05<br>F: -2.95 | Suppressed | Insulin-like growth factor binding protein 1; hepatic GH/IGF-1 axis regulator | Suppression may reflect impaired hepatic regulation of IGF and nutrient-responsive signaling (Chen et al., 2021; Leu & George, 2007), potentially compounding age-associated metabolic dysregulation. Low IGFBP1 is associated with insulin resistance and metabolic syndrome in humans (Kaushal et al., 2004). |
| *fkbp5*<br>(*LOC107385161*) | *FKBP5* | Male† | C: -3.20<br>F: -4.64 | Suppressed | FK506-binding protein 5; glucocorticoid receptor chaperone and stress response regulator | Suppression in both sexes suggests MC-LR-induced disruption of glucocorticoid-responsive stress signaling. FKBP5 regulates glucocorticoid receptor sensitivity and stress adaptation (Sato et al., 2025; Wang et al., 2018), potentially compromising an already age-sensitive stress-response system. |
| *stag3* | *STAG3* | Both† | M: +2.84<br>C: +2.18 | Induced | Stromal antigen 3; meiotic cohesin complex, chromosome segregation in germ cells | STAG3 is a meiosis-specific cohesin component required for chromosome segregation in germ cells, and pathogenic STAG3 variants are associated with premature ovarian insufficiency in humans (Demain et al., 2021). Its induction, together with increased RAD21L1-associated expression, suggests remodeling or engagement of cohesin-associated chromosome-maintenance programs following MC-LR exposure, potentially relevant to age-associated genomic vulnerability (Weeks et al., 2024). |

**Table 2:**
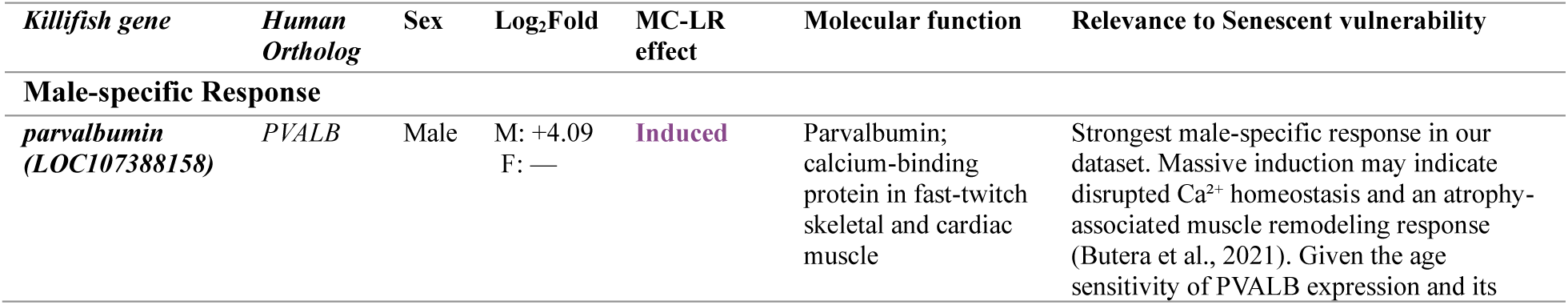
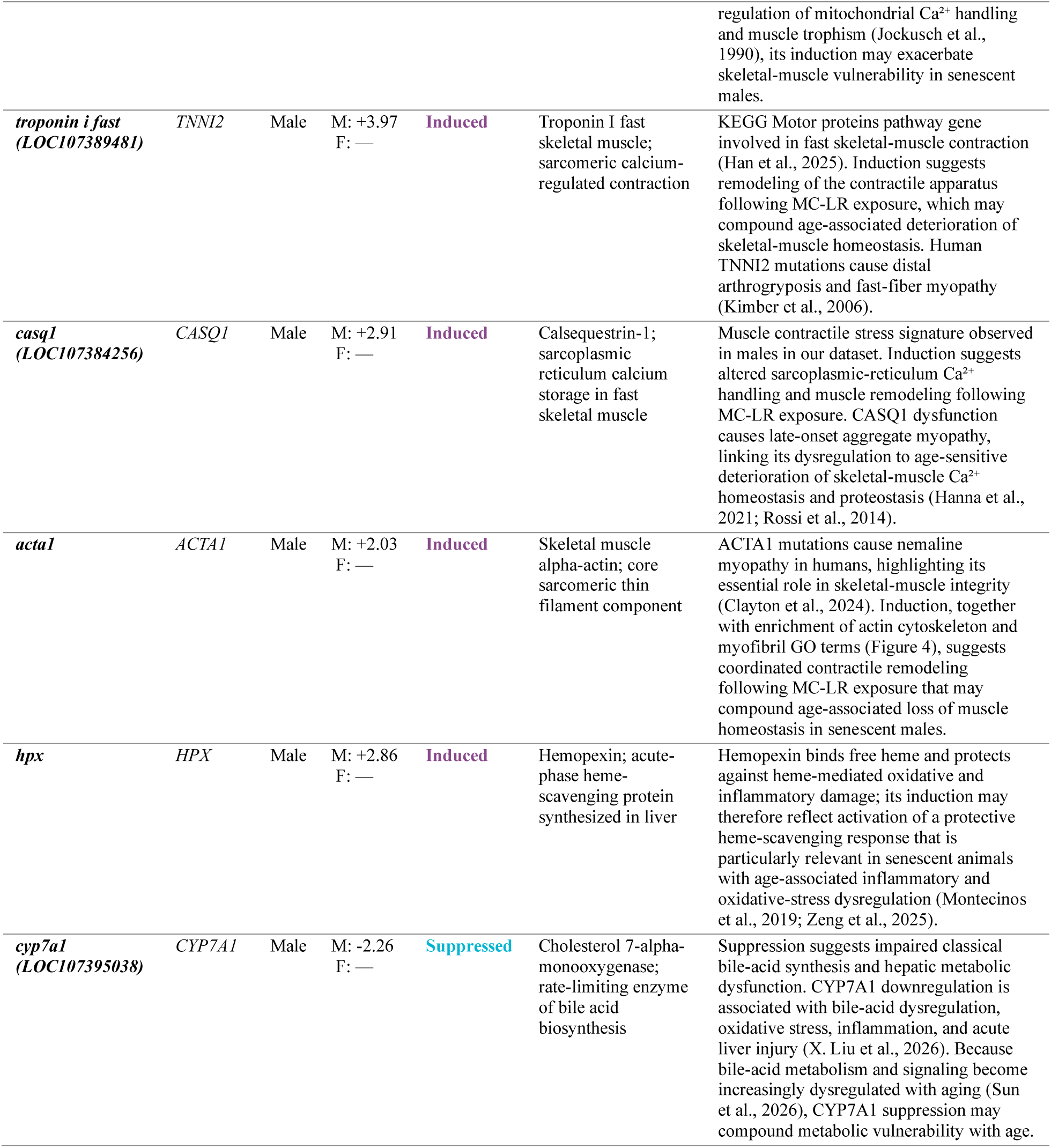
Selected differentially expressed genes in males; log₂FC, log₂ fold change (positive values indicate induction by MC-LR; negative values indicate suppression by MC-LR). Human orthologs were inferred from Swiss-Prot annotations (Ahmad et al., 2025) and represent putative functional homologs based on protein sequence similarity rather than experimentally validated orthology. M: reflects log2fold in males and F: reflects log2fold in females.

**Table 3:**
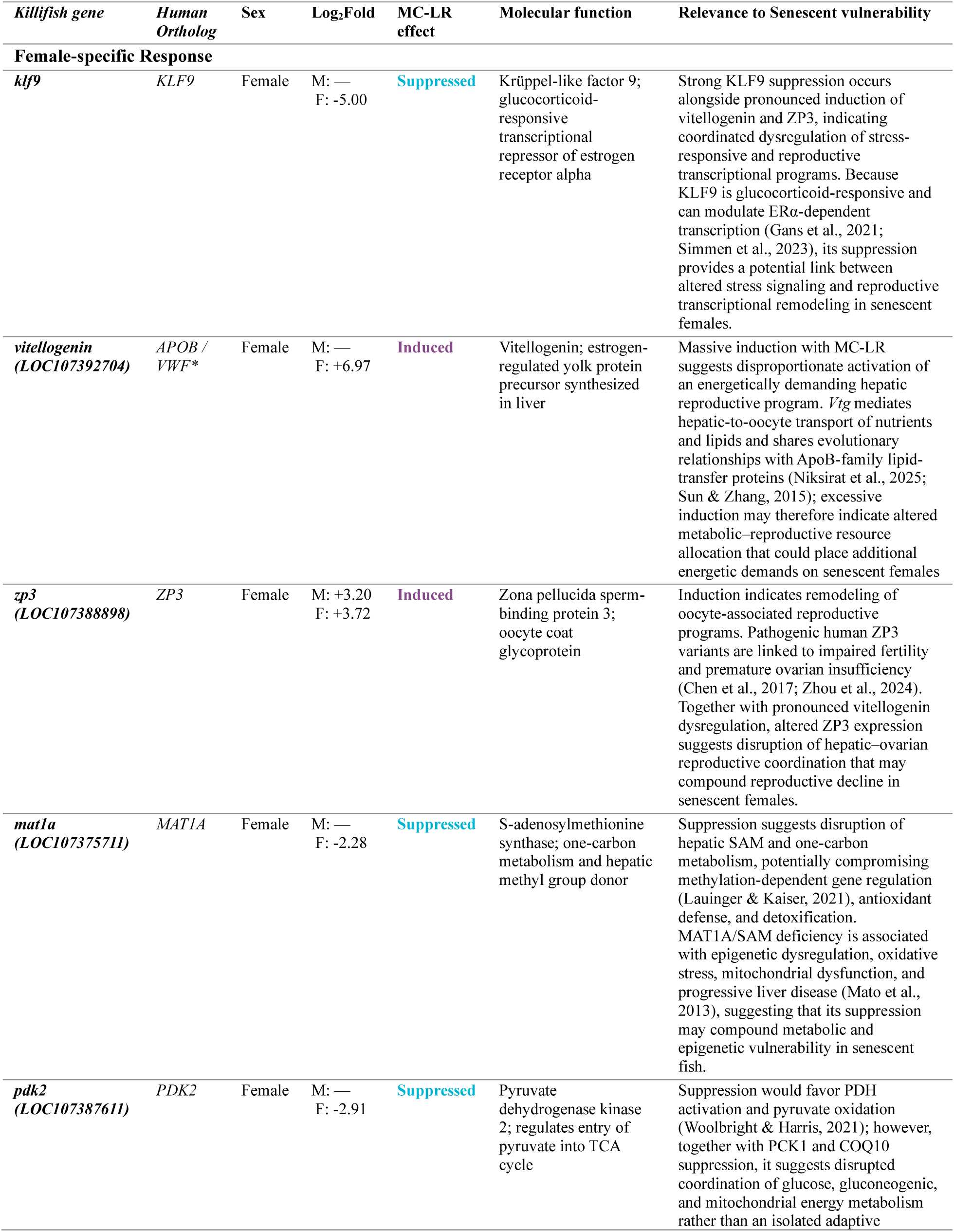
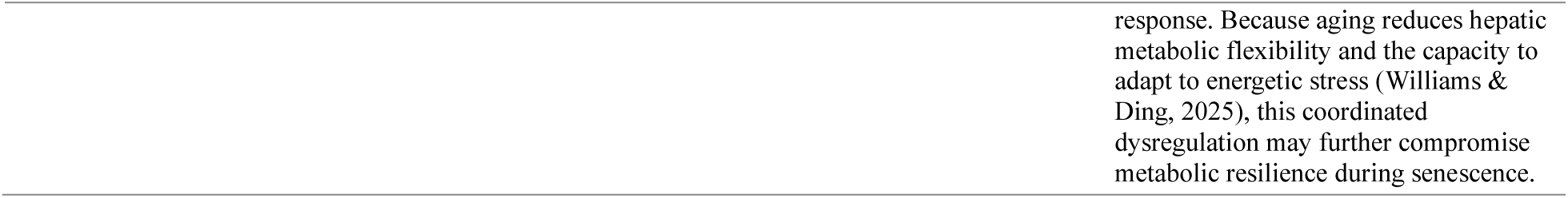
Selected differentially expressed genes in females; log₂FC, log₂ fold change (positive values indicate induction by MC-LR; negative values indicate suppression by MC-LR). Human orthologs were inferred from Swiss-Prot annotations (Ahmad et al., 2025) and represent putative functional homologs based on protein sequence similarity rather than experimentally validated orthology. *APOB/ VWF (Baker, 1988a, 1988b): Vitellogenin has no direct human ortholog; the indicated annotation reflects structural homology to human apolipoproteins. M: reflects log2fold in males and F: reflects log2fold in females.

Thus, substantially more transcriptional changes were detected when males and females were analyzed separately than when sexes were pooled. These findings indicate that the transcriptional response to MC-LR differs markedly between aged males and females and that analysis of the combined cohort obscures a substantial component of the exposure-associated response.

### A shared MC-LR-responsive gene set identifies common and divergent responses between sexes

Despite the broader sex-stratified responses, intersection of the male and female DEG sets identified 27 genes that were differentially expressed in both sexes, defining a core set of MC-LR-responsive genes (Fig. 3; Table 3). Of these, 22 showed concordant regulation between sexes, whereas five responded in opposite directions. Comparison of effect sizes further revealed substantial differences in response magnitude even among concordantly regulated genes (Fig. 3).

**Figure 3.**
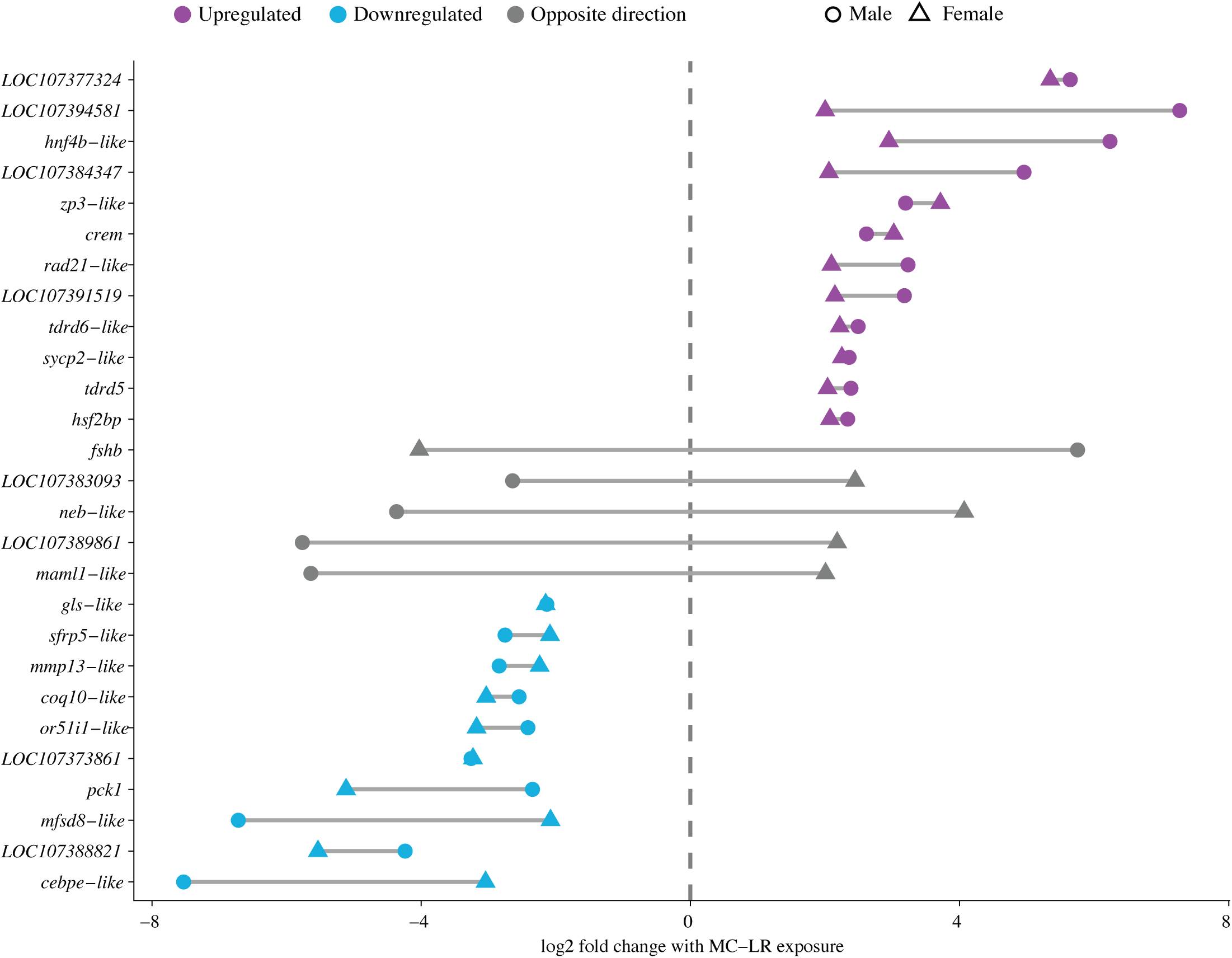
Sex-dependent regulation of shared MC-LR-responsive genes in senescent Nothobranchius furzeri. Connected-point plot comparing MC-LR-associated log₂ fold changes for the 27 genes identified as significantly differentially expressed in both male and female killifish (adjusted P ≤ 0.05 and |log₂ fold change| ≥ 2). Circles represent males and triangles represent females, with lines connecting the corresponding fold changes for each gene to illustrate differences in response magnitude and direction between sexes. Positive values indicate increased expression following MC-LR exposure and negative values indicate decreased expression. Purple indicates genes upregulated in both sexes, blue indicates genes downregulated in both sexes, and grey indicates genes exhibiting opposite directional responses between males and females. The vertical dashed line denotes no change (log₂ fold change = 0). The distance between connected male and female points illustrates the degree of sex-dependent divergence in the magnitude of the transcriptional response.

Several shared DEGs were associated with metabolic regulation. *Pck1*, encoding phosphoenolpyruvate carboxykinase 1, was strongly reduced following MC-LR exposure in both males and females. Reduced expression of *coq10-like* and *gls-like* accompanied the *pck1* response, identifying metabolic and mitochondrial-associated gene programs among the most prominent shared transcriptional changes. *Mmp13-like* was similarly reduced in both sexes.

A second group of shared genes showed increased expression following MC-LR exposure. These included *hnf4b-like*, *rad21-like* and *crem*, together with several genes associated with germ-cell and reproductive programs, including *tdrd5*, *tdrd6-like*, *sycp2-like* and *zp3-like* (Fig. 3). Thus, the concordant response encompassed metabolic, transcriptional, genome-regulatory and reproductive-associated genes.

Five shared DEGs showed opposing responses between sexes. Most notably, fshb increased in males but decreased in females. *Maml1-like*, *neb-like*, *LOC107389861* and *LOC107383093* likewise exhibited opposite directional responses (Fig. 3). The shared DEG set therefore represents a transcriptionally sensitive core rather than a uniform response to MC-LR, comprising both conserved responses and genes whose regulation remains strongly dependent on sex.

### MC-LR remodels global and tissue-associated transcriptional programs in a sex-dependent manner

The sex-dependent response was also evident at the whole-transcriptome level. Principal-component analysis of global gene expression separated samples according to both treatment and sex (Fig. 5A). PC1 accounted for 55.3% of total transcriptional variance and prominently separated untreated from MC-LR-exposed samples, whereas PC2 accounted for an additional 13.3% of variance and distinguished male and female transcriptional profiles. Biological replicates clustered closely within each sex-by-treatment group.

Functional enrichment analysis further revealed distinct biological programs underlying these responses. Analysis of the combined-sex DEG set identified enrichment of metabolic and catalytic functions, including carboxylic acid, organic acid, carbohydrate, monosaccharide, hexose and small-molecule metabolic processes, together with hydrolase, lyase, isomerase and tetrapyrrole-binding activities (Fig. 4A). Sex-stratified analyses revealed substantially broader and qualitatively distinct enrichment profiles (Fig. 4B,C, Supp Fig. 4), consistent with the greater number of DEGs detected when males and females were analyzed separately.

**Figure 4.**
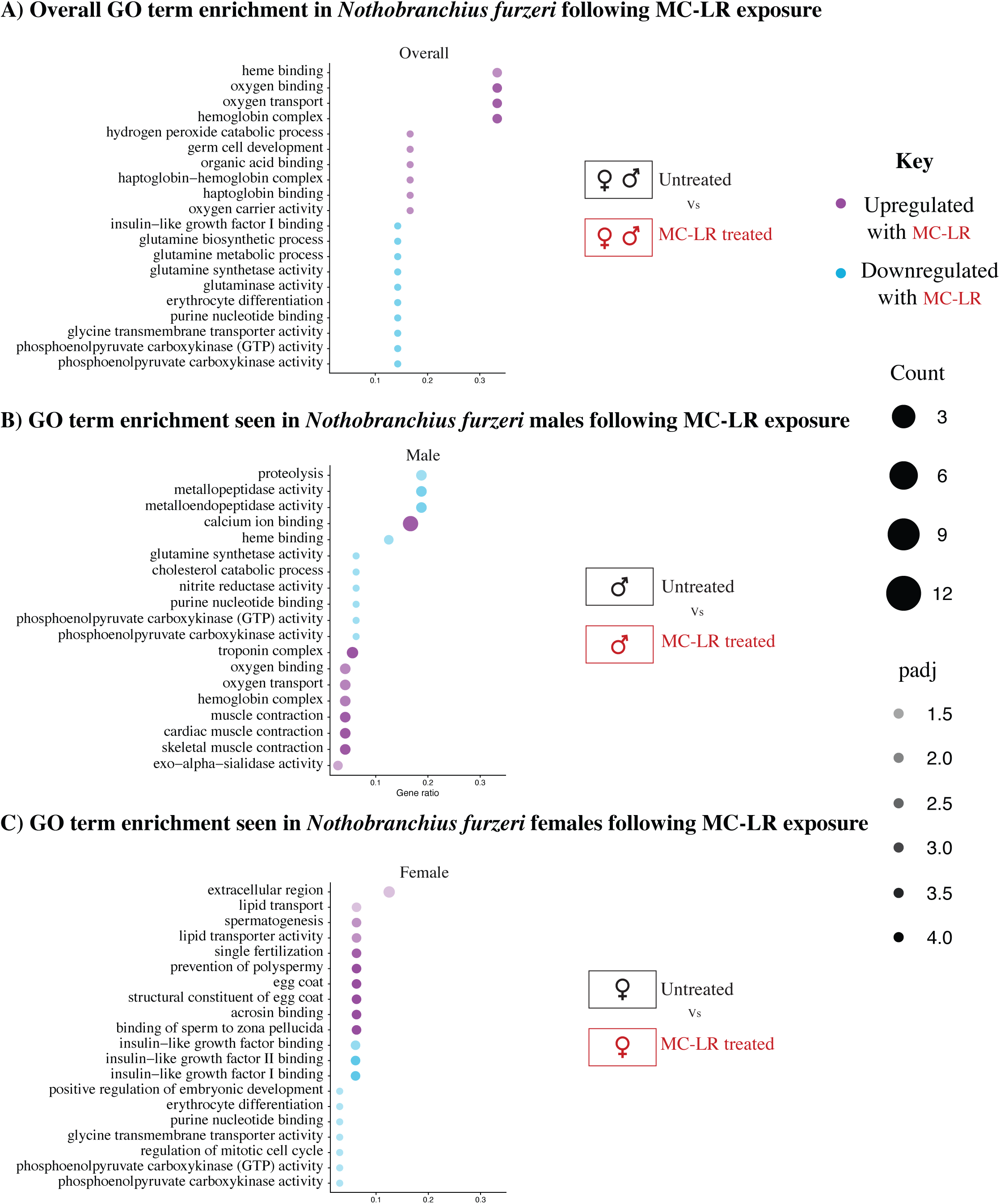
Gene Ontology enrichment analysis of transcriptional responses to MC-LR exposure in senescent Nothobranchius furzeri. Gene Ontology (GO) enrichment analysis was performed using differentially expressed genes identified following MC-LR exposure (adjusted P ≤ 0.05 and |log₂ fold change| ≥ 2). A) GO terms enriched in the combined-sex analysis. B) GO enrichment in males. C) GO enrichment in females. For each plot, the x-axis represents the Gene Ratio (the proportion of input genes associated with a given GO term), point size represents the number of genes contributing to the enriched term (Count), and point color represents either genes upregulated (magenta) and downregulated (cyan) and the intensity of color represents the adjusted P value (padj).

To further examine whether these transcriptional responses reflected coordinated changes in tissue-associated programs, we quantified marker-gene modules representing major cell and tissue signatures. MC-LR exposure produced distinct shifts in these signatures between males and females (Fig. 5B, Supp Fig. 5). Because these scores were derived from whole-body bulk RNA-seq, they represent changes in cell/tissue-associated transcriptional programs rather than direct measurements of cell abundance. Together, these analyses show that the relatively limited transcriptional response observed in the pooled cohort reflects the convergence of substantially different male and female responses to MC-LR.

**Figure 5.**
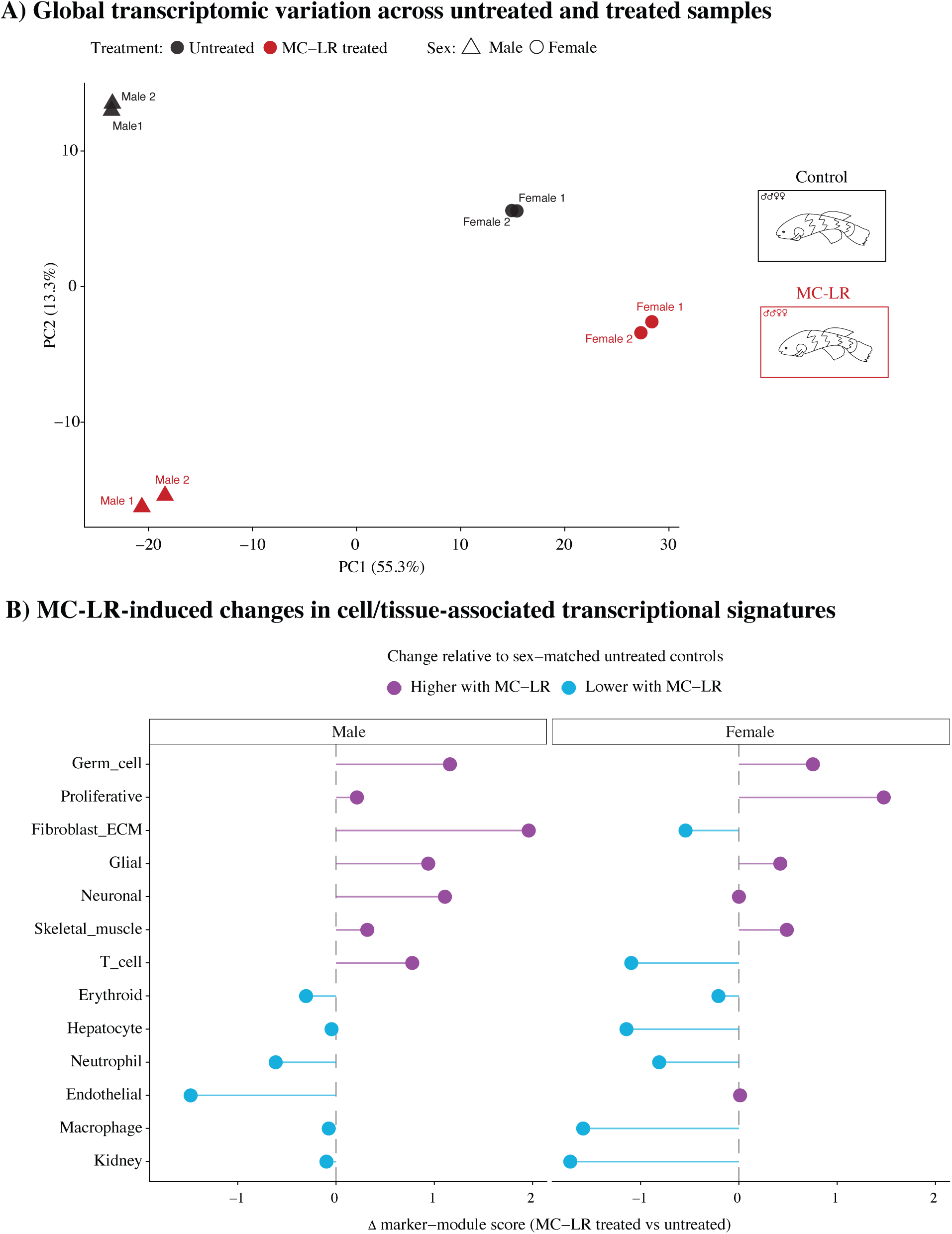
Global and cell/tissue-associated transcriptional responses to MC-LR exposure in senescent *Nothobranchius furzeri*. A) Principal-component analysis (PCA) of global gene expression across untreated and MC-LR-exposed male and female killifish. Circles represent females and triangles represent males; untreated samples are shown in black and MC-LR-exposed samples in red. B) Sex-stratified changes in cell/tissue-associated transcriptional signatures following MC-LR exposure. Marker-module scores were calculated from the expression of genes associated with each indicated cell/tissue signature, and changes are shown relative to sex-matched untreated controls (Δ marker-module score, MC-LR − untreated). Positive values (purple) indicate higher marker-module scores following MC-LR exposure, whereas negative values (blue) indicate lower scores. Marker-module scores represent coordinated expression of cell/tissue-associated gene sets and should not be interpreted as direct estimates of cell-type abundance.

### Male responses are enriched for contractile and extracellular-matrix-associated programs

The male transcriptional response was distinguished by prominent changes in genes associated with skeletal muscle structure, contraction and calcium handling. Differentially expressed genes included *pvalb*, *tnni2*, *casq1*, *acta1* and *myl3*, consistent with coordinated modulation of contractile machinery following MC-LR exposure.

Functional enrichment supported this pattern, with male DEGs enriched for muscle-associated functions, including oxygen binding and transport, hemoglobin complex, muscle contraction, cardiac muscle contraction, and skeletal muscle contraction (Fig. 4B). Other enriched terms included proteolytic, catalytic, and metabolic functions, reflecting additional components of the male transcriptional response.

Marker-module analysis provided an additional view of this response. Relative to sex-matched untreated controls, MC-LR-exposed males showed the largest positive shift in the fibroblast/extracellular-matrix-associated transcriptional signature (Fig. 5B). Germ-cell, neuronal, glial, T-cell and skeletal-muscle-associated signatures also increased, whereas endothelial, neutrophil and erythroid signatures decreased. Hepatocyte, macrophage and kidney-associated signatures showed comparatively modest changes. The concordance between gene-level differential expression, functional enrichment and marker-module analysis therefore identifies contractile and extracellular-matrix-associated transcriptional remodeling as prominent features of the male response to MC-LR.

### Female responses preferentially alter reproductive, metabolic and extracellular programs

Females exhibited a transcriptional response distinct from that observed in males. Among the strongly altered transcripts were genes associated with reproductive and liver-derived functions, including vitellogenin- and zona pellucida-associated genes. Female responses also included genes involved in metabolic and endocrine regulation, including *mat1a*, *pdk2* and *igfbp1*, indicating coordinated effects across reproductive and metabolic transcriptional programs.

Consistent with these gene-level responses, GO enrichment in females identified extracellular-region and lipid-transport terms, together with reproductive-associated functions and insulin/IGF-related binding activities (Fig. 4C). Additional enriched terms included glycine transmembrane transporter activity and regulation of mitotic cell cycle, further supporting a broad transcriptional response involving extracellular, reproductive, and metabolic functions.

Marker-module analysis further distinguished the female response from that of males. MC-LR exposure increased proliferative and germ-cell-associated transcriptional signatures, with smaller positive shifts in skeletal-muscle and glial signatures (Fig. 5B). In contrast, macrophage-, kidney-, hepatocyte-, neutrophil-, T-cell- and fibroblast/ECM-associated signatures decreased relative to untreated females.

Several of these changes were directionally distinct between sexes. Most notably, the fibroblast/ECM-associated signature showed the largest positive shift in males but decreased in females, while the T-cell-associated signature increased in males and decreased in females (Fig. 5B). These contrasting marker-module responses complement the sex-stratified differential-expression and functional-enrichment analyses and further demonstrate that males and females engage distinct transcriptional programs following MC-LR exposure.

## Discussion

Here, we define the transcriptional response to acute MC-LR exposure in naturally aged *Nothobranchius furzeri* and identify a strongly sex-dependent architecture of response. Although analysis of the pooled cohort revealed a relatively modest number of differentially expressed genes, stratification by sex uncovered substantially broader transcriptional changes in both males and females. This divergence was also evident at the level of global transcriptomic structure, functional enrichment, and cell/tissue-associated marker programs. Together, these findings indicate that biological sex is a major determinant of the response to MC-LR in aged animals and that analysis of males and females together can obscure biologically distinct responses to the same environmental stressor.

Despite this divergence, males and females shared a core set of MC-LR-responsive genes. Among the most prominent was *pck1*, which was strongly reduced in both sexes, with a greater magnitude of suppression in females. PCK1 catalyzes a key regulatory step in gluconeogenesis and contributes to metabolic adaptation during fasting and energetic stress. Its coordinated reduction, together with changes in *coq10-like*, *glul-like*, and other metabolic regulators, identifies energy and metabolic homeostasis as a common axis of vulnerability following MC-LR exposure. This response may be particularly relevant in aged animals, in which mitochondrial performance and metabolic flexibility are already altered. However, because these measurements were obtained from whole-body RNA-seq, determining whether the observed changes reflect altered hepatic glucose metabolism, systemic energetic stress, or coordinated responses across multiple tissues will require functional and tissue-resolved studies.

The shared DEG set also revealed that common sensitivity does not necessarily imply a common regulatory outcome. Although most shared genes changed concordantly between males and females, a subset exhibited opposing responses, including *fshb*, *maml1-like*, and *neb-like*. These directionally divergent responses reinforce the broader observation that the same exposure can engage distinct regulatory programs in males and females even when the same genes are transcriptionally sensitive. Such divergence may be particularly important in aged animals, where sex-dependent endocrine and metabolic states differ substantially from those earlier in life The male response was distinguished by coordinated changes in genes associated with muscle contraction and calcium handling. Differentially expressed genes included *pvalb*, *tnni2*, *casq1*, *acta1*, and *myl3*, accompanied by enrichment of muscle-associated functions, including muscle contraction, cardiac muscle contraction, and skeletal muscle contraction, as well as oxygen binding, oxygen transport, and hemoglobin complex. Additional enriched terms included proteolytic, catalytic, and metabolic functions. Marker-based transcriptional signature analysis similarly showed increases in skeletal muscle-associated and fibroblast/ECM-associated programs in males. Together, these analyses indicate that the transcriptional response to MC-LR in aged males encompasses contractile and oxygen-associated programs alongside broader metabolic and tissue-associated changes.

One possible explanation is that these changes reflect systemic adaptation to altered metabolic homeostasis. Skeletal muscle and liver are metabolically coupled through pathways including the glucose–alanine cycle, raising the possibility that disruption of hepatic metabolism could secondarily influence muscle-associated transcriptional programs. Alternatively, MC-LR may influence muscle-associated pathways independently of hepatic dysfunction. The present data cannot distinguish between altered tissue composition, compensatory transcriptional remodeling, and direct effects within skeletal muscle. Thus, rather than indicating muscle catabolism per se, the findings identify muscle-associated transcription as an important component of the systemic response to MC-LR in aged males.

Females exhibited a distinct response characterized by alterations in reproductive, metabolic, and extracellular transcriptional programs. Strong changes in vitellogenin- and zona pellucida-associated genes suggest perturbation of reproductive and endocrine-linked transcription, while alterations in *mat1a*, *pdk2*, *igfbp1*, and other metabolic regulators indicate concurrent changes in metabolic programs. GO enrichment further identified extracellular-region and lipid-transport terms, reproductive-associated functions, and insulin/IGF-related binding activities, together with additional metabolic and cell-cycle-associated functions. Marker-module analysis similarly identified increased proliferative and germ-cell-associated signatures together with reductions in hepatocyte-, macrophage-, kidney-, neutrophil-, and T-cell-associated programs. Collectively, these patterns suggest that the female response to MC-LR spans reproductive, extracellular, metabolic, and immune-associated transcriptional programs rather than being dominated by a single functional pathway.

Altered expression of stress- and endocrine-responsive regulators, including *klf9*, provides a potential link between these programs. KLF9 is responsive to glucocorticoid signaling and can influence endocrine transcriptional networks, raising the possibility that stress-axis activation contributes to the female response. However, the current data do not establish a direct glucocorticoid–KLF9–estrogen receptor mechanism. Measurements of hormone levels, receptor activity, and tissue-specific transcription will be necessary to determine whether this pathway directly mediates the reproductive-associated changes observed here.

The broader marker-module analysis reinforced the pronounced sex dependence of the response. Males showed increases in skeletal muscle-, neuronal-, glial-, T-cell-, and fibroblast/ECM-associated transcriptional signatures, whereas females exhibited a different pattern that included increased proliferative and germ-cell-associated signatures and reduced macrophage-, kidney-, hepatocyte-, neutrophil-, and T-cell-associated programs. Several signatures, including fibroblast/ECM and T-cell-associated programs, shifted in opposite directions between sexes.

Because these analyses were derived from whole-body bulk RNA-seq, they should not be interpreted as direct measurements of cell abundance. Rather, they identify coordinated changes in transcriptional programs associated with particular tissues or cellular states and provide a framework for future tissue-resolved and single-cell studies.

The aging context may be particularly important for interpreting these responses. Advanced age is accompanied by reduced metabolic flexibility, altered immune function, declining tissue repair capacity, and changes in endocrine regulation, all of which may modify how an organism responds to environmental stress. Our results therefore raise the possibility that MC-LR exposure interacts with pre-existing age-associated physiological states rather than simply eliciting a stronger version of the response seen in younger organisms. Direct comparison of young and aged animals will be necessary to determine which components of the response identified here are specific to aging and whether MC-LR accelerates or amplifies endogenous age-associated transcriptional programs.

More broadly, these findings argue that the biological consequences of cyanotoxin exposure cannot be fully understood without considering both age and sex. The simultaneous perturbation of metabolic, contractile, reproductive, immune, and extracellular transcriptional programs points to a systemic response whose organization differs markedly between males and females. Naturally aged *N. furzeri* therefore provides an experimentally tractable vertebrate model in which to examine how declining physiological reserve shapes responses to environmental toxicants across multiple biological systems.

### Limitations

Several limitations should guide interpretation of these findings. Most importantly, the study included two biological replicates per sex and treatment group. Although biological replicates clustered closely and several transcriptional responses were large in magnitude, the small sample size limits statistical power and the ability to estimate inter-individual variability. These findings should therefore be considered discovery-level and prioritized for independent validation in larger cohorts. Validation of prominent metabolic, contractile, and endocrine-responsive genes, including *pck1*, *hnf4*-associated genes, *mmp13*, *coq10*, and *klf9*, will be important for establishing the reproducibility of the observed responses.

A second limitation is the use of whole-body RNA-seq. This approach captures integrated organismal responses but cannot determine whether transcriptional changes arise from altered regulation within a specific tissue, changes in the relative representation of tissues or cell populations, or both. Accordingly, the cell/tissue-associated marker-module analysis should be interpreted as evidence of altered transcriptional signatures rather than changes in cell abundance. Tissue-specific RNA-seq, histological analyses, and single-cell or single-nucleus approaches will be required to localize these responses and determine their cellular basis.

Finally, although males and females exhibited markedly different transcriptional responses, formal sex-by-treatment interaction testing was not included in the present analysis. Terms such as sex-dependent or sex-specific should therefore be interpreted descriptively rather than as evidence of a statistically established interaction effect. Larger factorial studies will be required to directly test sex × treatment interactions. In addition, the GRZ strain, an inbred strain that has an exceptionally compressed lifespan and pronounced age-associated phenotypes, the extent to which these responses generalize to other *N. furzeri* strains or vertebrate species remains to be determined.

### Conclusions

Acute MC-LR exposure elicits a pronounced and sex-dependent transcriptional response in naturally aged *N. furzeri*. Although males and females share a core set of altered metabolic and regulatory genes, including strong suppression of *pck1*, their broader transcriptional responses diverge substantially. Males show prominent remodeling of contractile, skeletal muscle, extracellular-matrix, and neural-associated programs, whereas females display distinct changes in reproductive, metabolic, extracellular, and immune-associated transcription.

These findings position both aging and biological sex as important determinants of the response to cyanotoxin exposure. Rather than supporting a single tissue-specific mechanism, the transcriptomic data point to a systemic response involving interconnected metabolic and tissue-associated programs that are organized differently in males and females. More broadly, the study establishes naturally senescent *N. furzeri* as a tractable model for investigating how physiological aging shapes the response to environmental stress and provides a foundation for future tissue-resolved, functional, and age-comparative studies of MC-LR toxicity.

**Supp Figure 1** Hierarchical clustering of differentially expressed genes following MC-LR exposure in senescent *Nothobranchius furzeri*. Hierarchical clustering heatmap of the 36 significant differentially expressed genes (DEGs) identified in the combined-sex analysis of untreated (n = 4) and MC-LR-exposed (n = 4) GRZ killifish. Rows represent individual DEGs, and the color scale indicates log₂-normalized gene expression.

**Supp Figure 2** Hierarchical clustering of differentially expressed genes in male *Nothobranchius furzeri* following MC-LR exposure. Hierarchical clustering heatmap of the 313 significant differentially expressed genes (DEGs) identified between untreated male GRZ killifish and MC-LR-exposed males. Rows represent individual DEGs labeled by gene ID.

**Supp Figure 3** Hierarchical clustering of differentially expressed genes in female *Nothobranchius furzeri* following MC-LR exposure. Hierarchical clustering heatmap of the 263 significant differentially expressed genes (DEGs) identified between untreated female GRZ killifish and MC-LR-exposed females. Rows represent individual DEGs labeled by gene ID.

**Supp Figure 4** KEGG pathway enrichment analysis following MC-LR exposure in senescent *Nothobranchius furzeri*. KEGG pathway enrichment analysis was performed using differentially expressed genes identified following MC-LR exposure. A) Enriched KEGG pathways from the overall analysis including both sexes. B) Enriched pathways identified in the male-specific analysis. C) Enriched pathways identified in the female-specific analysis. Dot size represents the number of differentially expressed genes associated with each pathway (Count), and the x-axis indicates the GeneRatio, representing the proportion of input genes associated with each pathway. Dot color intensity represents the adjusted P value (padj), with lower values indicating greater statistical significance.

**Supp Figure 5** Expression patterns of cell- and tissue-associated marker genes following MC-LR exposure in senescent *Nothobranchius furzeri*. Hierarchical clustering heatmap showing normalized expression of representative marker genes associated with major cell and tissue transcriptional signatures across untreated and MC-LR-exposed male and female *N. furzeri.* Columns represent individual biological replicates, with treatment and sex indicated by annotation bars above the heatmap. Rows represent marker genes grouped according to their associated cell or tissue signature. Expression values were scaled across samples for each gene, with warmer colors indicating relatively higher expression and cooler colors indicating relatively lower expression. Hierarchical clustering of marker genes illustrates distinct expression patterns across sex and treatment groups and was used to derive the cell/tissue-associated transcriptional signatures.

## Author Contributions

Z.A.: Investigation, Methodology, Formal analysis, Writing – original draft. C.H.: Methodology. D.K.: Conceptualization, Methodology, Supervision, Funding acquisition, Writing – review and editing, Corresponding author.

## Ethics Statement

All animal experiments were conducted under Protocol No. DK10112024 approved by IACUC at North Carolina Central University.

## Data Availability

All data will be deposited on the GEO database and available to download.

## Acknowledgments

We acknowledge the Julius L. Chambers Biomedical/Biotechnology Research Institute (BBRI) at North Carolina Central University for providing research facilities and resources. We thank the BBRI faculty and staff and students for their support in this project, Vandana Veershetty, Evan E. Pittman, Dr. Derek Norford, Ms. Camilla Felton, Thomas J Broek, Dr. Qing Cheng, and Dr. Claudia Alberico. We are grateful to Dr. Anne Brunet and members of the Brunet Laboratory at Stanford University for their guidance in establishing the killifish facility and colony. We also thank Riley Galton and Robert Schnittker from the Sánchez Alvarado Laboratory at the Stowers Institute for all their support and expertise in guiding us for killifish embryo and juvenile husbandry, and Wouter Lanneau of Microbiotests for his assistance with killifish embryos. We also acknowledge the use of an AI agent for discussion and interpretation of the study findings.

Use of AI was done in July and August 2026 to assist with editing, refinement, organization and clarity of manuscript text, and development and troubleshooting of R code used for data visualization. It was not used to generate any of the main analysis or an underlying experimental data. All AI-assisted content, code, analyses, and interpretations were done according to established guidelines (Flanagin et al., 2026) and independently reviewed and verified by the authors, who take full responsibility for the accuracy and integrity of the manuscript.

