## Supplementary material for "Low-Dose Microcystin-LR Elicits Sex-Dimorphic Transcriptomic Responses in Senescent *Nothobranchius furzeri*: Implications for Cyanotoxin Vulnerability in Aging Vertebrates": Supp Figures

**Supplementary Figure 1**

**Hierarchical clustering heatmap for overall comparison (both sexes; 36 DEGs)**

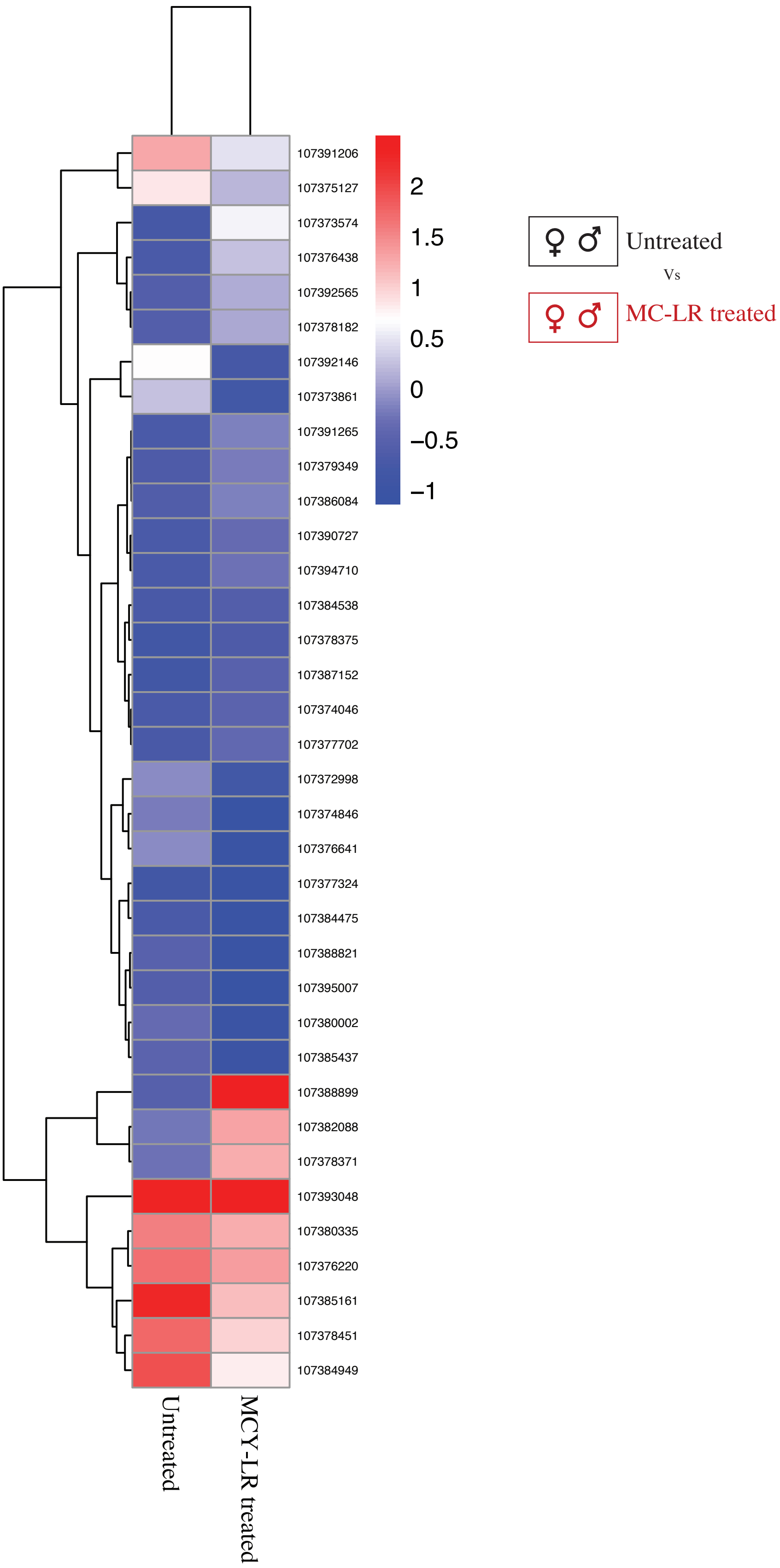

### Supplementary Figure 2

### Hierarchical clustering heatmap for males (313 DEGs)

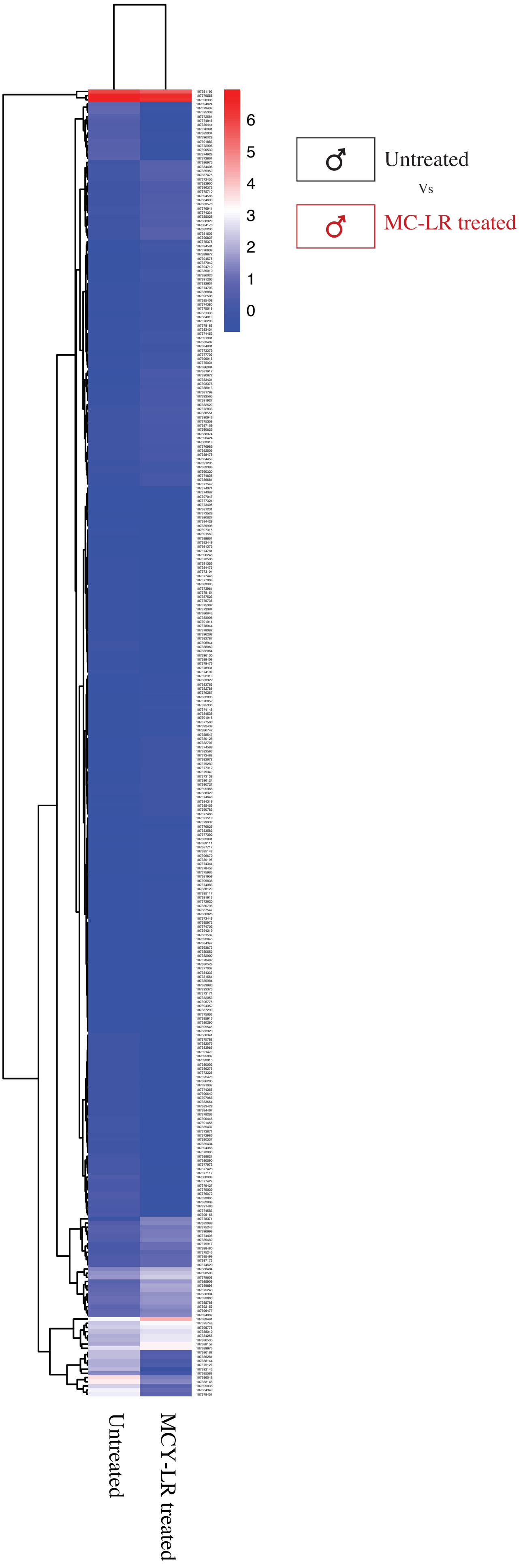

**Supplementary Figure 3**

**Hierarchical clustering heatmap for females (263 DEGs)**

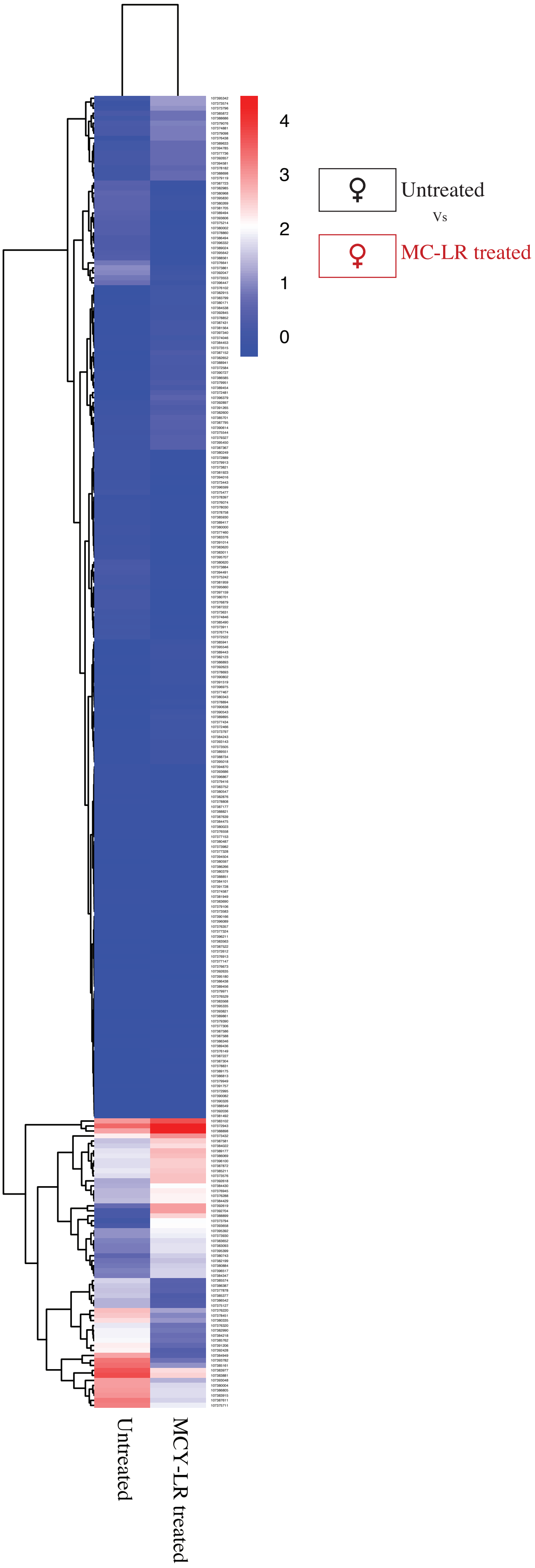

**Supplementary Figure 4**

**A) Overall KEGG analysis in *Nothobranchius furzeri* following MC-LR exposure**

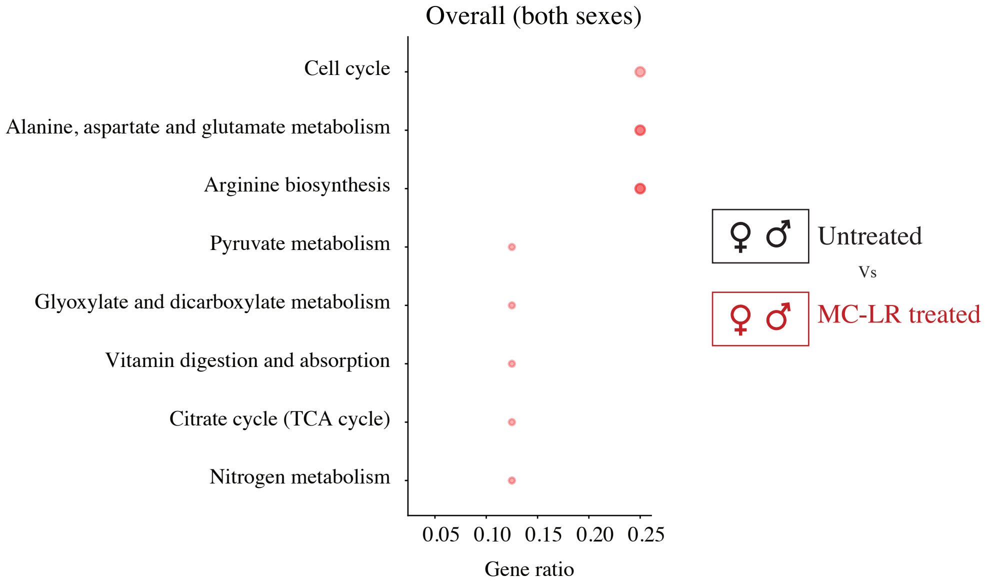


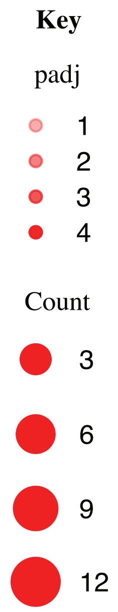


**B) KEGG analysis in *Nothobranchius furzeri* males following MC-LR exposure**

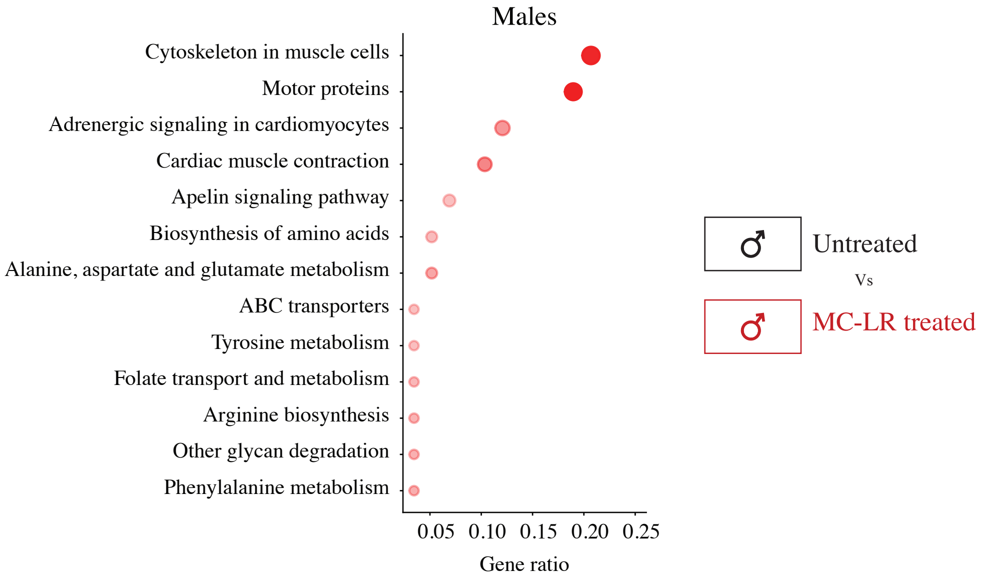


**C) KEGG analysis in *Nothobranchius furzeri* females following MC-LR exposure**

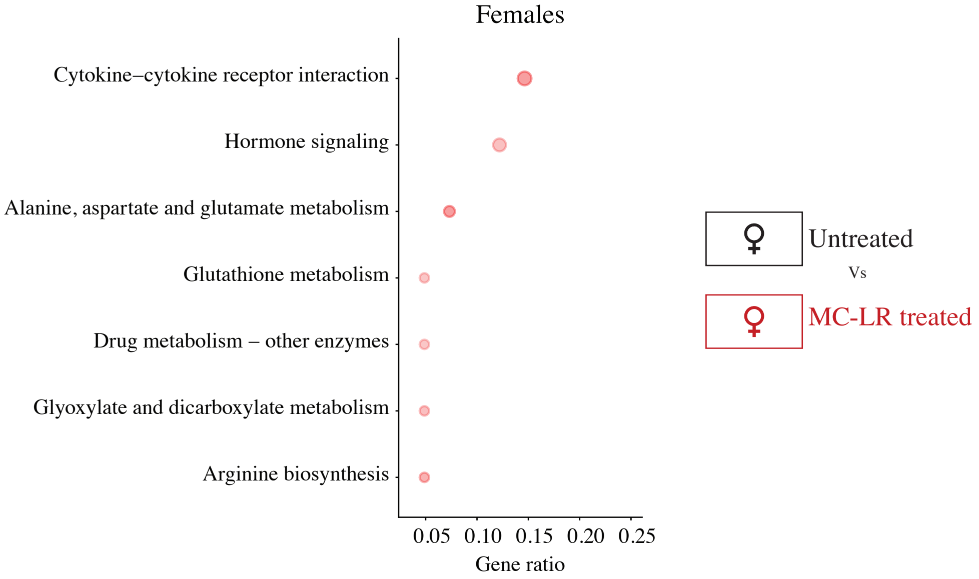

### Supplementary Figure 5

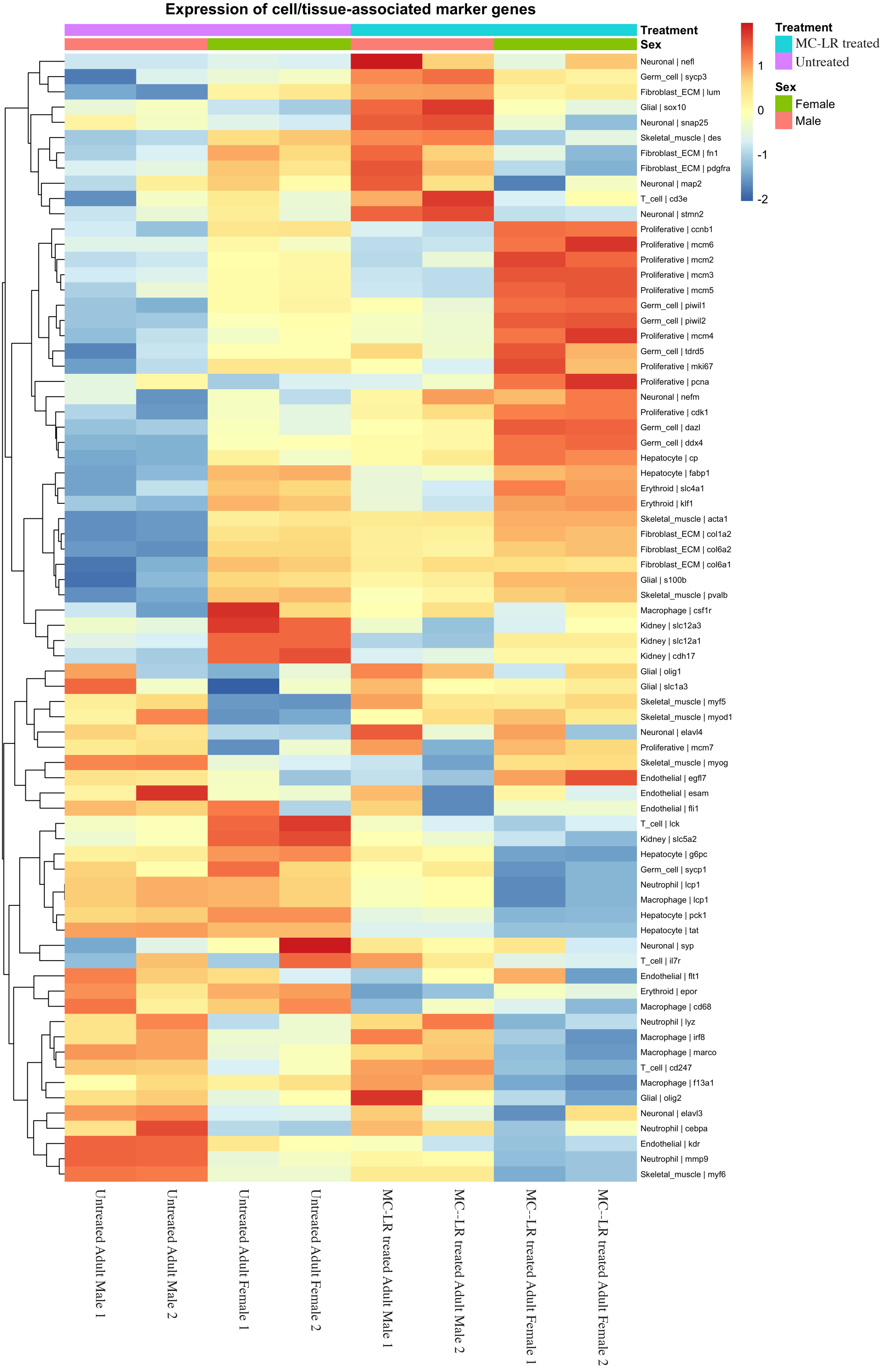
